# Limitations of lymphoblastoid cell lines for establishing genetic reference datasets in the immunoglobulin loci

**DOI:** 10.1101/2021.07.15.452448

**Authors:** Oscar L. Rodriguez, Andrew J. Sharp, Corey T. Watson

## Abstract

Lymphoblastoid cell lines (LCLs) have been critical to establishing genetic resources for biomedical science. They have been used extensively to study human genetic diversity, genome function, and inform the development of tools and methodologies for augmenting disease genetics research. While the validity of variant callsets from LCLs has been demonstrated for most of the genome, previous work has shown that DNA extracted from LCLs is modified by V(D)J recombination within the immunoglobulin (IG) loci, regions that harbor antibody genes critical to immune system function. However, the impacts of V(D)J on data generated from LCLs has not been extensively investigated. In this study, we used LCL-derived short read sequencing data from the 1000 Genomes Project (n=2,504) to identify signatures of V(D)J recombination. Our analyses revealed sample-level impacts of V(D)J recombination that varied depending on the degree of inferred monoclonality. We showed that V(D)J associated somatic deletions impacted genotyping accuracy, leading to adulterated population-level estimates of allele frequency and linkage disequilibrium. These findings illuminate limitations of using LCLs for building genetic resources in the IG loci, with implications for interpreting previous disease association studies in these regions.

**Author summary:** Lymphoblastoid cell lines (LCLs) are cells that have been manipulated to proliferate indefinitely in order to provide a replenishable source of DNA. However, because these cell lines are derived from B cells which have undergone V(D)J recombination they contain somatic deletions within regions of the genome that encode antibody genes. Although several large collaborative projects have utilized DNA from LCLs to generate invaluable genomic resources for the scientific community, the negative impacts of cell line artifacts in these regions of the genome have not been fully appreciated. In this study, we used newly released sequencing data from a large collection of LCLs to determine that the non-inherited artificial deletions within the antibody gene loci can have detrimental effects on downstream genetic analyses.

## Introduction

Lymphoid blastoid cell lines (LCL) are generated by infecting B cells with the Epstein Barr Virus (EBV)[1] to create immortalized cell lines. Various consortia, including The International HapMap Project[2, 3], 1000 Human Genome Project (1KGP)[4–6], Genome In A Bottle[7, 8] and Human Genome Structural Variation Consortium[9] have used DNA from LCLs to characterize common genetic variation, generate gold standard sets of small insertions and deletions (indels), and comprehensively genotype structural variants (SV). Variant call sets from these initiatives have been instrumental to the genomics community, and are routinely used in genome-wide association studies (GWAS) and other genetic studies. Genome-wide genotypes from LCLs have been shown to be nearly identical to genotypes derived from whole blood or peripheral blood mononuclear cells (PBMC) using SNP arrays[10], whole exome sequencing[11, 12] and whole genome sequencing[13]. However, somatic LCL-associated alterations are present in particular regions of the genome, namely within the immunoglobulin (IG) heavy (IGH) and light (lambda, IGL; kappa, IGK) chain loci. These alterations could impact sequencing, mapping, and genotype results in these regions, with potential implications for downstream uses of these data.

The IG loci encode the variable (V), diversity (D) and joining (J) gene segments that serve as the building blocks for the expression of functional B cell receptors (BCRs) and antibodies (Abs). During B cell development, the V, D, and J gene segments within each IG locus (V and J in the case of IGL and IGK) are somatically rearranged through a process called V(D)J recombination[14]. During this process, intervening DNA between recombined V, D, and J segments is excised. The size of these somatic deletions on the recombined chromosome depends on the selected V, D, and J genes, but can extend 100’s of Kb, and will vary from cell to cell. Collectively, DNA isolated across a pool of B cells (e.g., naive B cells) representing many independent V(D)J recombination events would be expected to represent each germline haplotype present in a given sample (Fig. 1). In contrast, a pool of B cells originating from a single or dominant expanded B cell would harbor DNA not fully representative of both paternal and maternal germline haplotypes within the IG loci (Fig. 1). In the latter instance, genotyping methods dependent on different read alignment signatures such as read depth/coverage, discordant read mapping, soft-clipped or split reads could produce inaccurate germline genotypes.

**Figure 1.**
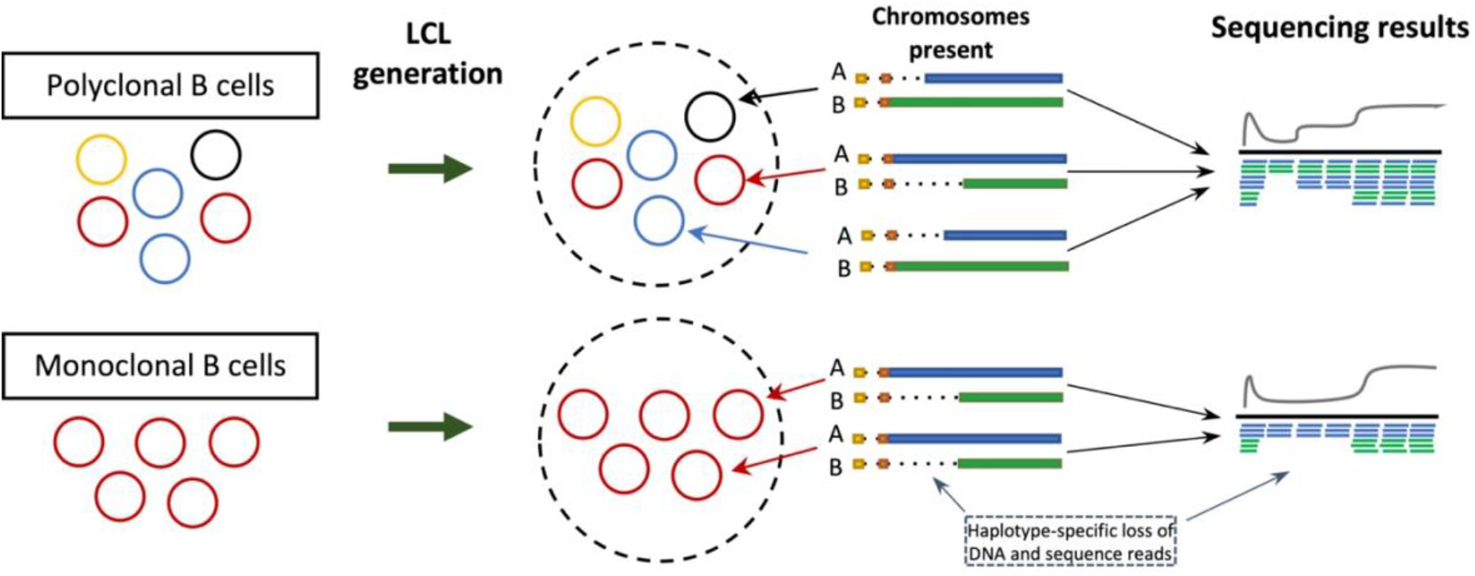
Generation of LCLs can lead to loss of sequencing data. LCLs are generated from a pool of B cells. A pool of polyclonal B cells will contain different V(D)J recombination events and therefore the collection of different B cell clones would lead to no loss of sequencing data from either haplotype. LCLs generated from monoclonal B cells, *i.e.* B cells with the same single V(D)J event, would lead to loss haplotype-specific sequencing data.

Recent long read sequencing and assembly of complete IGH haplotypes from selected 1KGP individuals has revealed the presence of V(D)J recombination associated deletions[15], indicating that genotypes derived from such samples within regions impacted by V(D)J recombination are inaccurate. While it has been speculated previously that V(D)J recombination would have negative impacts on LCL-derived sequencing data [16–18], this has not been comprehensively investigated. Given this, we sought to evaluate the extent of sample-level V(D)J recombination in LCL-derived short read sequencing data from the 1KGP, and assess downstream impacts of these somatic events. We demonstrate that short read data is affected by V(D)J recombination and, depending on the sample, is derived from either single dominant or multiple B cell clones. We show that variation in sample clonality is associated with variability in genotyping accuracy, negatively impacting estimates of allele frequency and linkage disequilibrium (LD). These data raise important considerations for using 1KGP genotypes to augment genetic association studies in the IG loci, and in addition to other issues discussed previously[16–18], may further explain the paucity of disease associations within these complex regions of the genome.

## Materials/Subjects and Methods

### 1000 Human Genome Project Data

Paired-end 150 bp PCR-free 30X coverage Illumina data on 2504 individuals from the 1KGP[19] was downloaded from the European Bioinformatics Institute (EBI) under the study ID ERP114329. 1KGP phase 3[4] SNPs were downloaded from ftp://ftp.1000genomes.ebi.ac.uk/vol1/ftp/data_collections/1000_genomes_project/release/20190312_biallelic_SNV_and_INDEL/ALL.chr14.shapeit2_integrated_snvindels_v2a_27022019.GRCh38.phased.vcf.gz.

### Assessment of insert sizes, read depth, and the identification of V(D)J recombination events

Insert sizes for read pairs within each sample were calculated using the Picard (https://github.com/broadinstitute/picard) CollectInsertSizeMetrics tool with the following parameters: `--DEVIATIONS 1000000 --MINIMUM_PCT 0` for reads spanning chr14:105862198-107043718. The same tool and parameters were used for ten random 1.2 MB windows across the genome. To calculate read depth across the IGHD and V regions, we used samtools[23] (IGHD, hg38, chr14:105,865,458-105,939,756; IGHV, hg38, chr14:105,939,756-106,883,718). To identify read pairs within each sample representing V(D)J recombination events, we utilized a custom python script. For each sample, the number of clones was calculated by counting the number of unique IGHV and IGHJ gene pairs detected. The frequency of each “clone” was calculated by determining the number of reads mapping to a unique IGHV and IGHJ gene pair, and taking this as a fraction of the total number of reads assigned to any IGHV/IGHJ pair. The sizes of somatic deletions were determined by calculating the genomic distances between the IGHJ and IGHV genes utilized in a given V(D)J event.

### Analysis of heterozygosity, allele frequency, and LD

The percentage of heterozygous SNPs was calculated for each sample in the centromeric and telomeric region of the selected V gene in the dominant clone. Two VCFs were created for each region using tabix to select the region of interest on the 1KGP phase 3 VCF. The telomeric region was always set to start at the beginning of *IGHV6-1* (chr14:105,939,756; GRCh38). The fraction of heterozygous SNPs was calculated by counting the number of heterozygous SNPs over the total number of SNPs in each VCF.

To assess the effects of V(D)J recombination on allele frequency estimates and LD, we subsetted the 1KGP phase 3 genotype call set to samples from the “AFR” superpopulation, further selecting samples representing two extremes of clonality (0-25%, n=38; 75-100%, n=38). Allele frequencies for each set of samples was calculated using the vcftools `freq` tool. The LD scores were calculated for the African samples of each set of samples using the vcftools `hap-r2` tool with the parameters `--ld-window 1000000 --min-r2 0.01`.

## Results

### Detecting signatures of V(D)J recombination

To determine the effect of V(D)J recombination on whole genome sequencing (WGS) data in IGH, we used paired-end 150 bp PCR-free Illumina data on 2,504 individuals from the 1KGP, recently resequenced to high coverage[19]. The occurrence of V(D)J recombination results in large somatic deletions within the IGH locus spanning the IGHJ, IGHD, and IGHV regions. To assess signatures of these somatic deletions we first analyzed paired-end mapping distances, as measured by the predicted “insert sizes”. We reasoned that the presence of V(D)J recombination events would result in larger insert sizes, and that these would be enriched within IGH. To assess this, we calculated the number of read pairs with an insert size >900 bp (two times the library DNA insert size) at 10 random 1.2 MB windows (the length of the IGH locus in GRCh38) across the genome from five individuals chosen at random. Across these regions in the selected individuals, we observed that 0.08% to 0.12% (mean = 0.10%) of the paired-end reads contained an insert size greater than 900 bps. In contrast, across all samples, 0.13% to 1.55% (mean = 0.49%) of paired-end reads in IGH contained an insert size greater than 900 bps. This is almost a 5-fold increase in the number of paired-end reads with a larger insert size (Fig. 2A).

**Figure 2.**
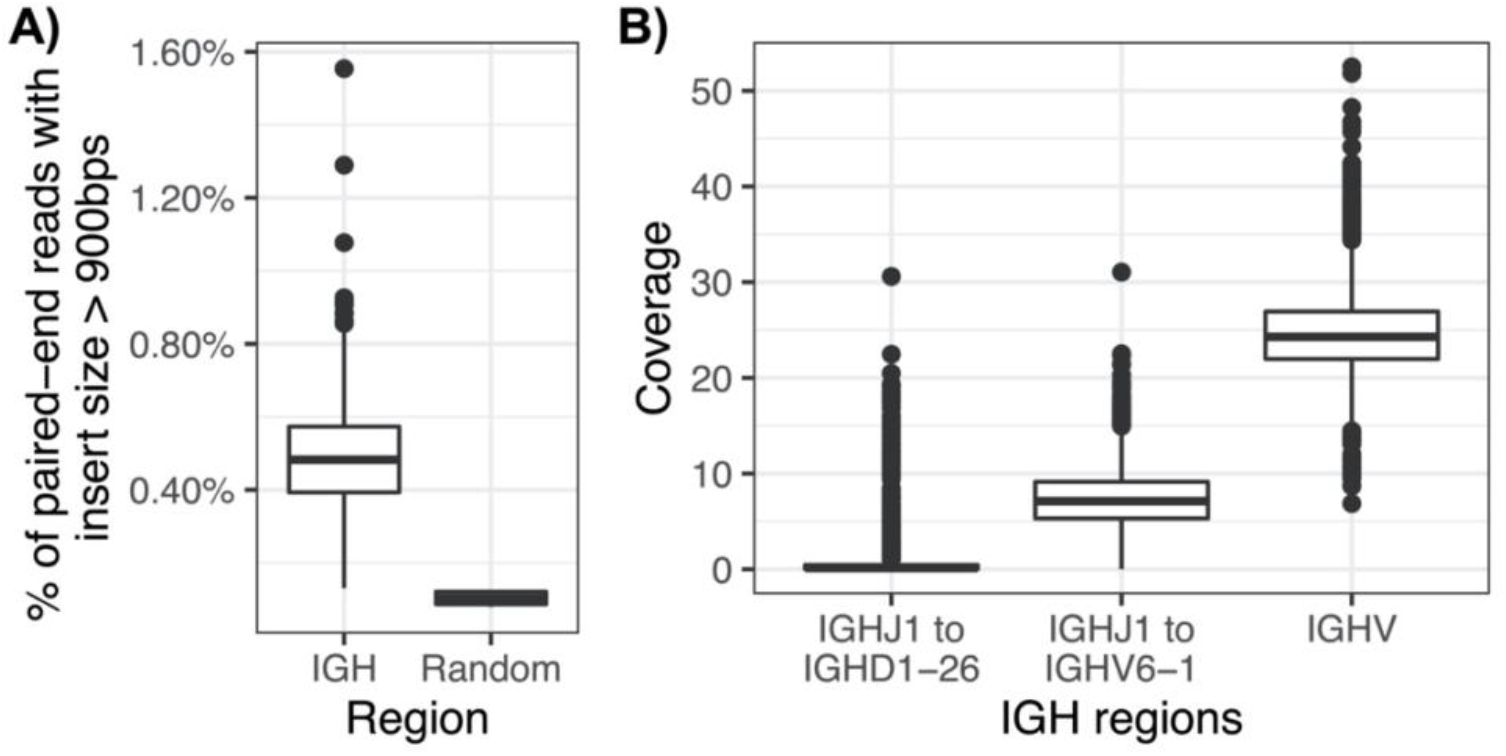
Signatures of V(D)J recombination in paired-end WGS data. (A) Percentage of paired-end read pairs with insert size greater than 900 bps in IGH and random genome-wide 1 Mb windows. (B) WGS coverage between (1) *IGHJ1*, the IGHJ gene closest to the telomeric end and *IGHD1-26*, the second closest gene to the IGHJ gene cluster, (2) *IGHJ1* and *IGHV6-1*, the IGHV gene closest to the IGHD and IGHJ gene cluster and (3) IGHV region

To further evaluate the effect of V(D)J recombination, we calculated the coverage over the IGHD region. During B cell development, through the formation of the pre-B cell receptor, V(D)J recombination results in a loss of DNA between selected IGHJ and IGHD gene segments on each homologous chromosome[20]. Therefore, if V(D)J recombination has occurred, there should be limited to no coverage within the IGHD region. The closest IGHJ and IGHD gene pair, *IGHJ1* and *IGHD7-27*, are within 100 bps, so we assessed coverage between *IGHJ1* and *IGHD1-26* (~15 Kb) across all 2504 individuals, and observed a mean coverage of 0.76X (range = 0 - 30.6). We also observed a mean coverage of 7.30X between *IGHJ1* and *IGHV6-1*, in contrast to a mean coverage of 24.68X across the entirety of the IGHV region (Fig. 2B).

We next used the paired-end data to directly detect V(D)J recombination events by identifying read pairs with one mate overlapping an IGHJ gene and the other mate overlapping an IGHV gene. We detected read pairs representing V(D)J recombinants in all 2,504 samples, with a mean of 22 V(D)J associated read pairs per sample (range = 2 - 66). V(D)J rearrangements utilizing *IGHJ4* and *IGHV3-23* were the most common (Supplemental Fig. 1). Taken together, these three pieces of evidence, increased insert sizes, decreased coverage over IGHD, and the direct detection of read pairs overlapping V(D)J recombination events, indicate that in fact LCLs utilized by the 1KGP cohort have undergone V(D)J recombination.

### Sequencing data derived from multiple B cell clones

Given that V(D)J recombination has occurred across the cohort, we sought to determine whether all samples were affected equally. We reasoned that samples with sequencing data from a single B cell clone (monoclonal) or from multiple clones (polyclonal) will be differentially impacted by the effects of V(D)J (Fig. 3A). We therefore sought to determine the number and frequency of V(D)Js in each sample. To do this, we assigned read pairs overlapping V(D)J events to their respective combination of IGHJ and IGHV genes. Reads across a given dataset harboring the same IGHJ/IGHV combination were grouped, and used as a proxy for a group of clonally related sequences. We thus took the number of grouped sequence reads to represent the frequency of a particular IGHJ/IGHV combination, which we heretofore refer to as a clone (Fig. 3A). Following this, we calculated the number of unique IGHJ/IGHV combinations (“clones”) present in each sample, and their relative frequency, allowing us to approximate the number of different B cell clones represented in a sample. We found that sequences across samples in the cohort were derived from a mean of 10.76 B cell clones (range = 1 - 30; Fig. 3B). From this, 18 samples were predicted to be monoclonal, represented by reads mapping to only a single clone. We also reasoned that polyclonal samples, represented by many clones, but in which the majority of sequencing data is predicted to be derived from a dominant clone will have profiles similar to those observed for monoclonal samples. To estimate this, we asked what proportion of all sequences containing a V(D)J recombination event were represented by the most frequently observed clone. In doing this, we found that in 407 and 88 samples, respectively, 50% and 75% of all reads containing V(D)J recombination events mapped to a single clone. This indicated that although these samples had a polyclonal signature, the majority of sequencing data was likely derived from a single dominant clone. Based on this approximation, across samples, the top clone identified contributed on average 32.94% (range = 5 - 100%) of the sequencing data (Fig. 3C). We observed a modest population bias, with African and European individuals containing a slightly greater fraction of sequencing data from a single clone (Supplemental Fig. 2).

**Figure 3.**
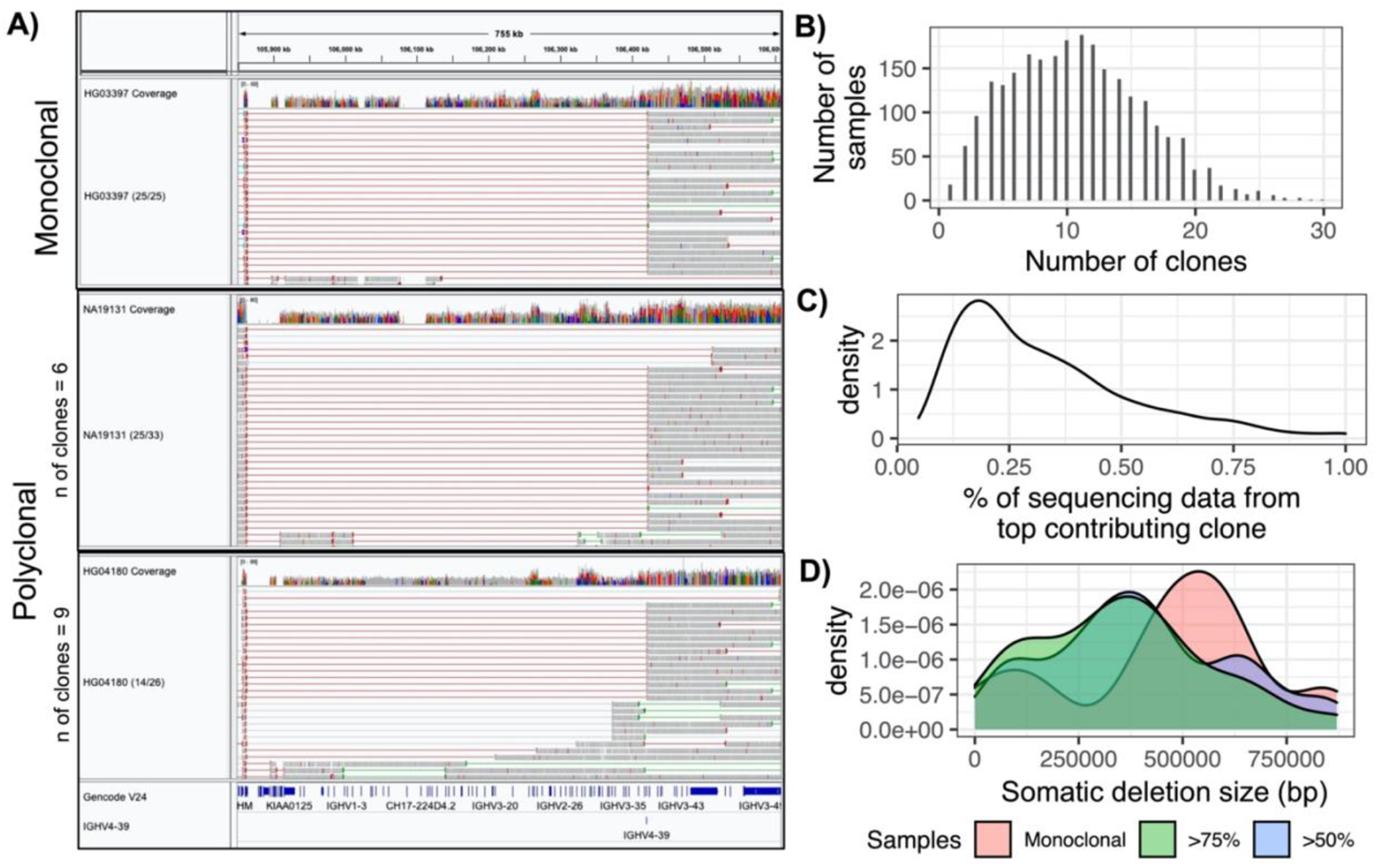
Clonality in different samples. (A) IGV screenshots showing three samples with different clonality but have a dominant clone with *IGHV4-39*. Red and gray lines represent large insert lengths. HG03397 has 25 read pairs aligning with to an IGHJ and IGHV gene, all of which align to *IGHV4-39* and hence labelled as monoclonal. NA19131 and HG04180, which are polyclonal, have 25/33 and 14/26 read pairs aligning to *IGHV4-39*. (B) Number of samples with different clonalities (C) Percentage of sequencing data derived from dominant clone (D) Amount of IGHV DNA lost in monoclonal samples and polyclonal samples with 75% and 50% of their sequencing data derived from the dominant clone

Depending on the clonality of the sample and the genes involved in the primary V(D)J recombination event present within a sample, the size of the region impacted was expected to vary. We estimated this in each sample based on the most prevalent IGHJ/IGHV gene combination observed, revealing that in many samples the predicted size of these somatic deletions was extensive. In the 18 monoclonal samples, we found that an average of 464 Kb of the IGH locus was impacted by V(D)J recombination (range = 74.3 - 937.2 Kb). In samples that were polyclonal, but still represented by a dominant clone (i.e., those in which >50% and >75% of sequence data were derived from a single IGHJ/IGHV combination), the regions impacted by V(D)J recombination were found to be on average 356 Kb (50%, range = 74.3 - 945.0 Kb) and 401 Kb (75%, range = 74.3 - 945.0 Kb) in size (Fig. 3D). Regions affected by V(D)J recombination in each sample are provided in Supplementary Table 1.

### The effects of V(D)J recombination on genotype call sets

We reasoned that V(D)J events could impact the accuracy of sample- and population-level genotypes in two primary ways: 1) the loss of DNA and reduced read coverage over extended regions of the locus would result in the increased likelihood of calling homozygous genotypes at heterozygous positions; 2) somatic hypermutations (SHMs) in recombined genes could introduce false heterozygous SNPs. Furthermore, these effects would likely be more prominent in monoclonal samples, as well as polyclonal samples represented by dominant clones.

To investigate these potential impacts, we first evaluated the number of heterozygous SNPs in the centromeric and telomeric regions of the recombined IGHV gene in monoclonal samples. We found that the mean percentage of heterozygous variants telomeric of the IGHV gene used for V(D)J recombination was 3.4 fold higher than the mean percentage of heterozygous variants centromeric of the IGHV gene (*P =* 0.003, two-sided paired Wilcoxon test; Fig. 4A). To assess this effect in polyclonal samples, we split individuals into four groups representing varying degrees of clonal bias, based on whether 0-25%, 25%-50%, 50%-75% and 75%-100% of sequencing data within a given sample came from the dominant clone. In samples with 0 to 25% of sequencing data from the dominant clone, for which we expected to observe minimal impacts on genotyping, the mean percentage of heterozygous variants telomeric (62%) of the selected recombined IGHV gene was 1.04-fold higher than in the centromeric region (59%; *P*=0.006, two-sided paired Wilcoxon test; Fig. 4B). We noted significant differences in the remaining three groups as well (Fig. 4B), but the average fold-differences between heterozygous percentages telomeric to the V(D)J event relative to centromeric to the V(D)J event were greater, and was greatest in samples from the 75%-100% group (2.08-fold).

**Figure 4.**
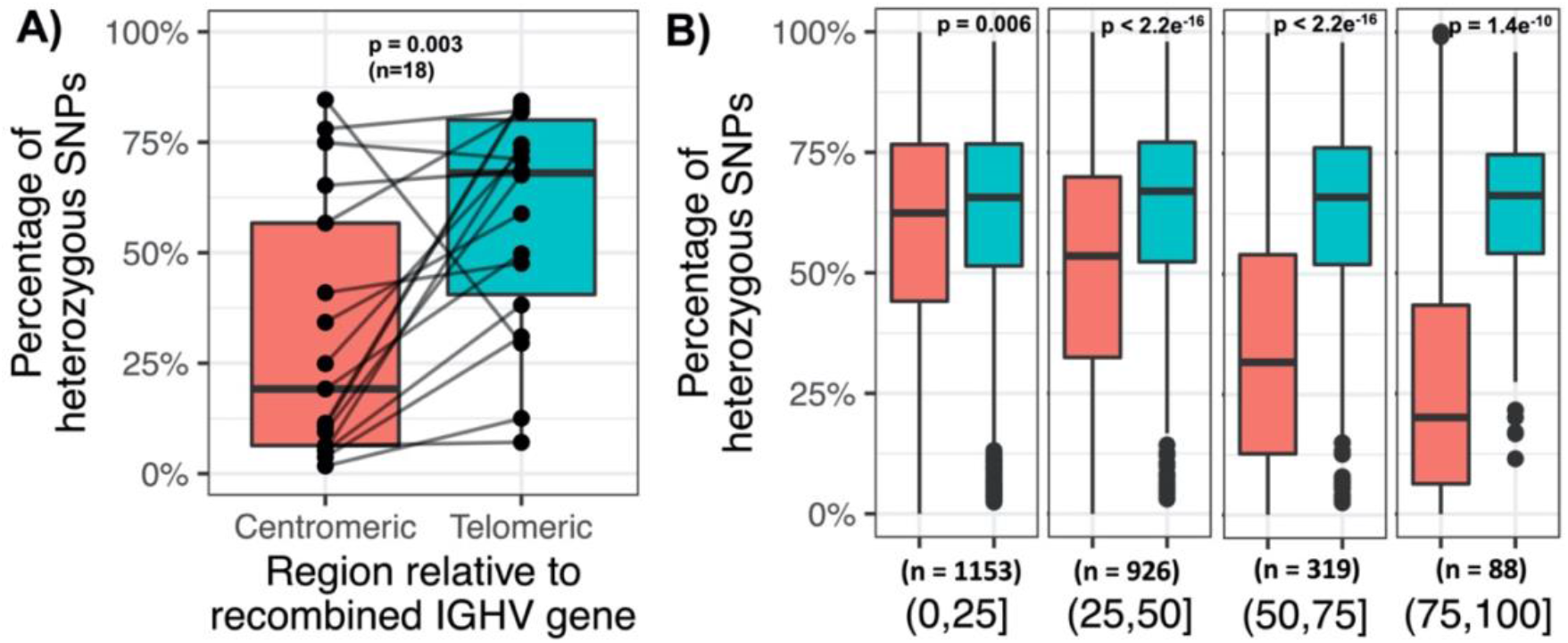
Difference in heterozygosity in centromeric and telomeric regions of selected V(D)J recombination IGHV gene and clonality bias. (A) Eighteen samples were predicted to be monoclonal. The percentage of heterozygous SNPs in the centromeric and telomeric regions of the IGHV gene selected for V(D)J recombination was calculated. Each line represents a single sample connecting the percentage of heterozygous SNPs in the centromeric and telomeric regions. (B) Samples were split based on their clonality bias. Samples in the “(0,25]” group, the least clonal group, derived up to 25% of sequencing data from the dominant clone and samples in the “(75,100]” group, most clonal group, derived 75% to less than 100% of their sequencing data from a dominant clone. Similar to (A), the proportion of heterozygous SNPs was calculated in the centromeric and telomeric regions of the IGHV gene selected for V(D)J recombination.

Additionally, we evaluated the number of heterozygous SNPs overlapping IGHV genes most frequently selected for V(D)J recombination in each sample. When stimulated by an antigen, B cells acquire SHMs within IG V, D, and J genes as a means to increase antibody affinity[21]. Therefore, SHMs in the recombined IGHV gene are more likely to be detected in monoclonal or polyclonal samples with reads primarily derived from a dominant clone. Indeed, there was a significant positive correlation (R = 0.33, p-value < 2.2e-16) between the contribution of sequencing data from the dominant clone and the number of the heterozygous variants within the IGHV gene selected by V(D)J recombination (Supplementary Fig. 3A). We also directly compared the number of heterozygous genotypes within the recombined IGHV genes to non-recombined IGHV genes across all samples, and observed an average of 1.92 heterozygous positions in the recombined IGHV genes, compared to 0.5 in the non-recombined IGHV genes (Supplementary Fig. 3B).

### The effects of V(D)J recombination on estimates of allele frequency and linkage disequilibrium

The previous section detailed the effects on genotypes due to V(D)J recombination. Given that genotypes are used to determine allele frequencies in a population, we set out to test if allele frequencies differed between samples that are more or less monoclonal. The allele frequencies of common SNPs (MAF > 0.05) were compared between samples within superpopulations with 0-25% (less monoclonal) and 75-100% (more monoclonal) of sequencing data derived from the dominant clone. Of the 4,354 SNPs analyzed, 1,258 (29%) had an allele frequency difference greater than 0.05 (Fig. 5A). Since V(D)J recombination excises DNA 3’ of selected IGHV genes, we would expect to observe more genotyping errors caused by V(D)J related somatic deletions within the centromeric region of the IGHV locus. Consistent with this, we observed greater differences in allele frequencies within the proximal (centromeric) region of the locus when comparing estimates generated from less monoclonal samples to more monoclonal samples (Fig. 5B).

**Figure 5.**
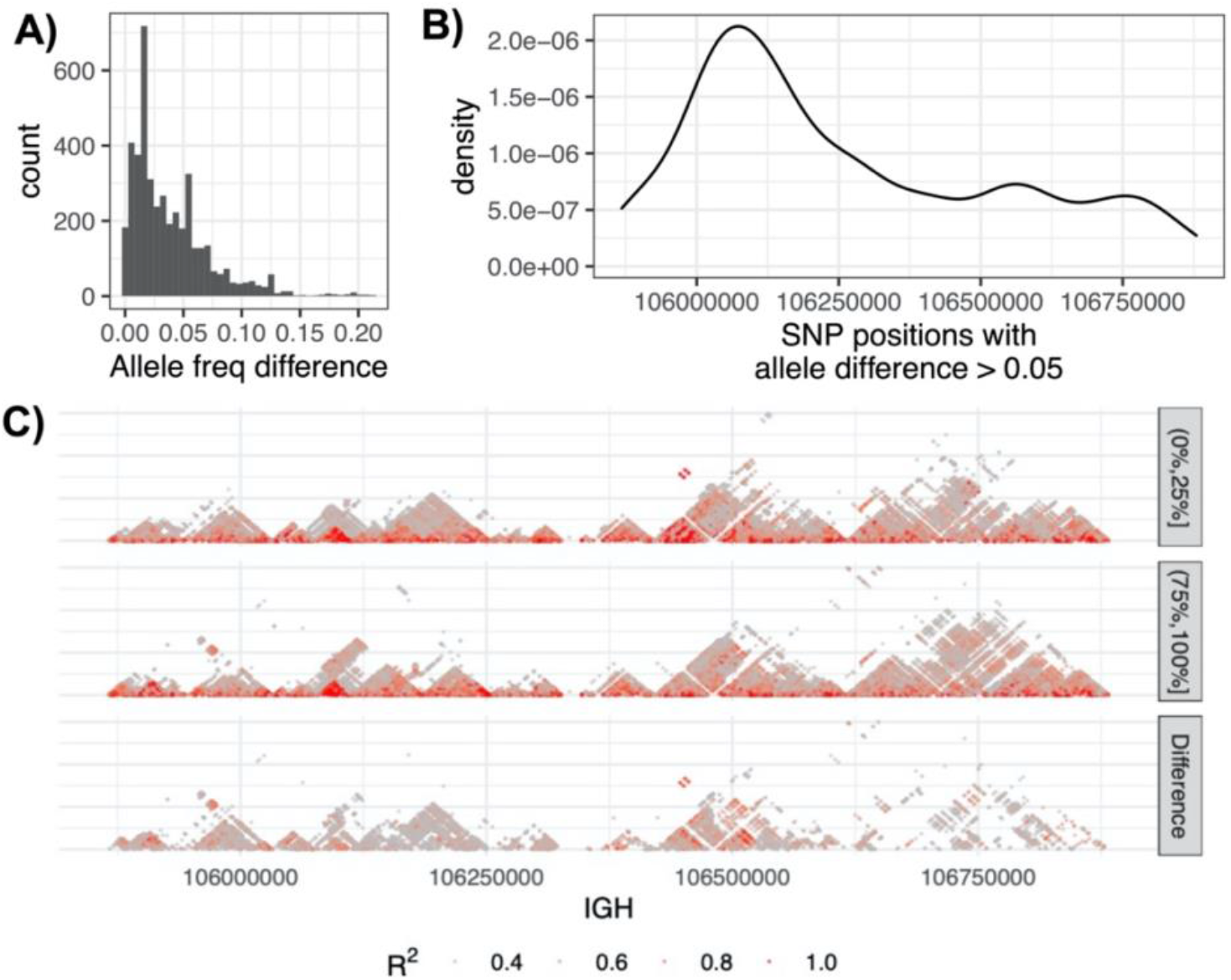
Allele frequency and LD differences in individuals with low and high clonality. (A) The distribution of common allele frequency difference between individuals with low clonality and high clonality, defined as 0 to 25% and 75% up to 100% of sequencing data was derived from a single clone, respectively. (B) Position in IGH with common allele frequency differences greater than 0.05 (C) LD for African individuals with low (“(0%,25%]”) and high clonality (“(75%,100%]”), and the difference in LD between both groups.

Genotypes are also used for the calculation of LD between SNPs. Given the demonstrated impact on allele frequency estimates, we reasoned that effects on genotype accuracy would also impact LD estimates. To assess this, we chose 76 samples from the African superpopulation representing extremes of clonality. Two groups of 38 samples each from the lower clonality group and the higher clonality group were selected. The LD *r*^2^ values across the locus were computed and compared between the two groups (Fig. 5C), revealing different LD structure. We found that 11% (236,827) of the SNP pairs exhibited differences in LD (*r*^2^) greater than 0.1 (Supplementary Fig. 4). The differences in allele frequencies and LD estimates observed here indicated that inaccurate genotypes resulting from impacts of V(D)J recombination also affect downstream analyses.

## Discussion

Previous studies have concluded that there are minimal differences between genotypes from matched LCL and non-LCL samples[10–13]. While true on a genome-wide scale, here we show that the impact on the IGH locus is more apparent due to V(D)J recombination. Using 1KGP samples (n=2504) recently resequenced on the Illumina NovaSeq platform to 30x coverage using PCR-free 2×150 bp libraries, we evaluated different sequencing features affected by V(D)J recombination and SHM within the IGH locus. Specifically, we demonstrated that signatures of V(D)J recombination within LCL-derived DNA can be observed, including increased insert sizes of read mate-pairs, decreased read coverage over the IGHD and proximal IGHV gene regions, as well as direct evidence of somatically recombined IGHJ and IGHV genes. By assessing the frequency of specific IGHJ/IGHV recombination events within each sample, we were able to estimate the number of approximate B cell clones likely represented within a sample, and determine the proportion of sequencing data derived from each B cell clone, revealing variation in clonality across samples. Importantly, we were able to determine that V(D)J recombination can result in loss of DNA spanning large segments of the locus, with clear impacts on variant genotyping. The extent of these effects varied between samples based on the gene segments involved in the primary V(D)J recombination event, and the degree of monoclonality observed. Together these observations highlight critical limitations of using LCLs to develop comprehensive reference resources for the IGH locus at the sample and population level.

It has previously been argued that the locus complexity of IGH has made it difficult to study using high-throughput approaches such as short-read data and genotyping arrays[15–17]. This has impeded our ability to accurately characterize genetic diversity within IGH, and robustly test hypotheses about the functional role of IGH germline variation in disease risk and antibody-mediated immunity. The analyses we have conducted here indicate that the large-scale use of LCLs for establishing genetic reference panels in IGH may also present additional barriers to effectively interrogating IGH in genetic studies with downstream implications that need to be considered. For example, LCL-derived datasets such as the 1KGP have been critical for establishing population-genetic metrics across the genome, and have been used to augment GWAS and inform functional and population genetic studies. For example, consortia efforts such as gnomAD[22] have aggregated data from multiple sources, including LCL-derived data from the 1KGP, to power such studies. However, we have shown here that genotype, allele frequency, and LD estimates are incorrect for much of IGH due in part to impacts of V(D)J events in the data. This highlights a need to reconsider use of these cohorts for such purposes. We argue that, at a minimum, the continued use of LCL-derived datasets could be improved by removing erroneous genotypes caused by V(D)J recombination induced deletions. As part of this study, we have released a BED file with the coordinates of V(D)J recombined induced deletions for each sample (Supplementary Table 1). It is possible that the development of genotyping pipelines that account for such data anomalies on a per-sample basis would lead to more accurate estimates of genotype and allele frequencies within IGH, with potential downstream implications for improving imputation approaches utilized by GWAS. Finally, while the focus of our study has been on the IGH locus, these observations would be applicable to the IGL and IGK loci as well.

## Acknowledgements

This work was supported, in part, by grants from the U.S. National Institutes of Health NIH R24AI138963 (to CTW) and NIH 1F31NS108797 (to OLR). This work was supported in part through the computational resources and staff expertise provided by the Scientific Computing at the Icahn School of Medicine at Mount Sinai.

## Supplementary Figures

**Supp Fig 1.**
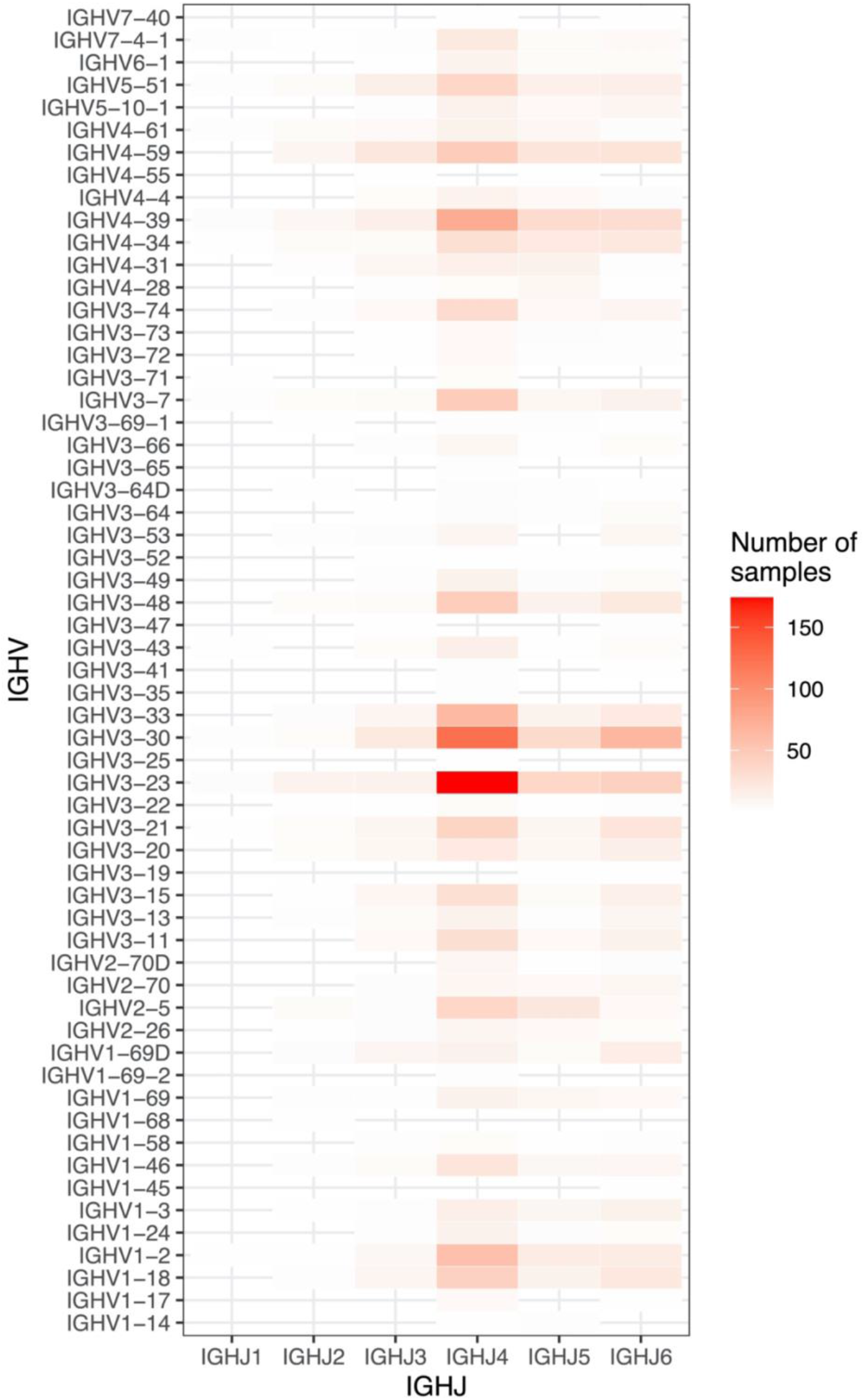
Number of samples with dominant clone containing IGHV and J pair.

**Supp Fig 2.**
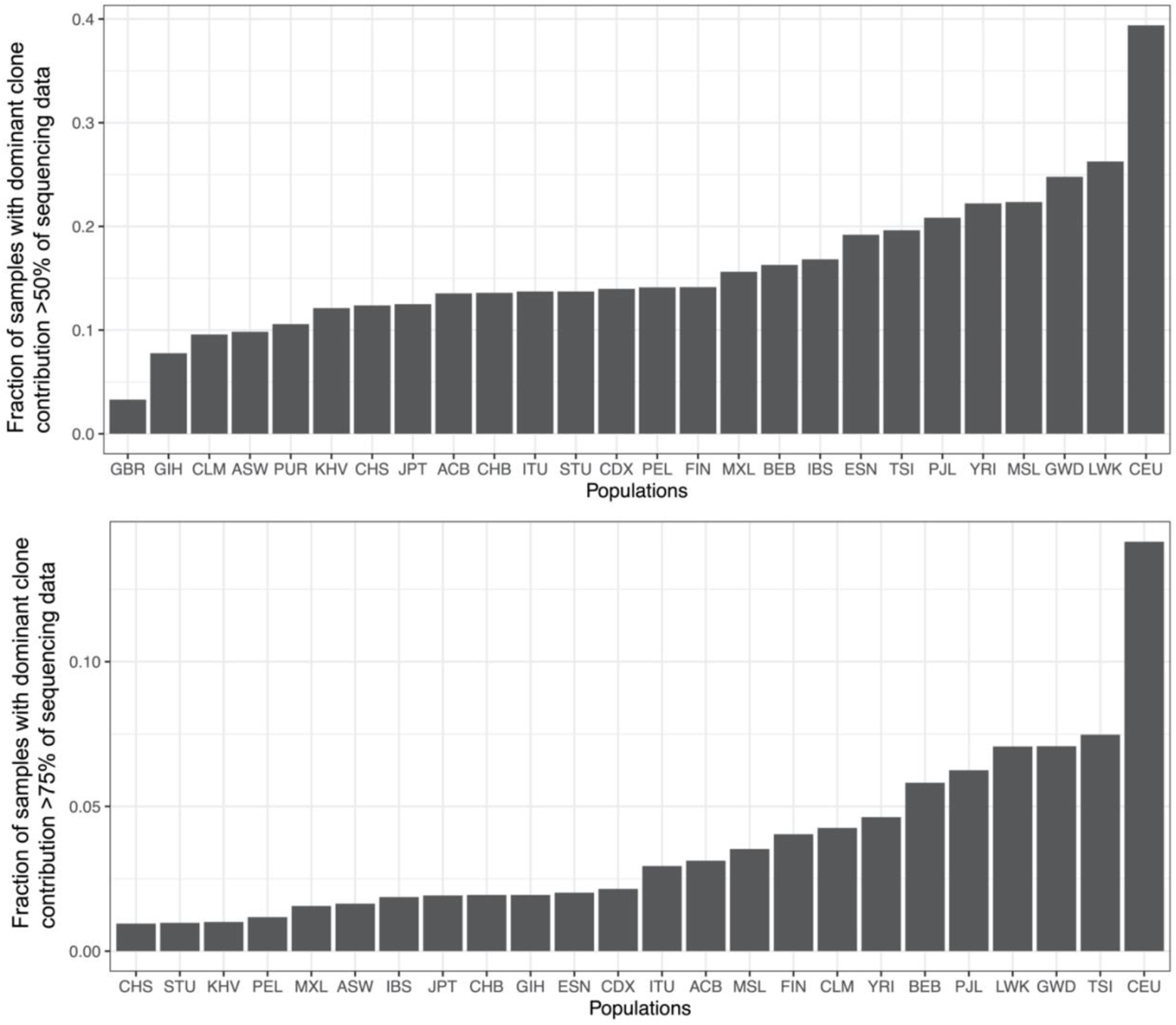
Fraction of samples per population where the dominant clone is more than 50% and 75% prevalent.

**Supp Fig 3.**
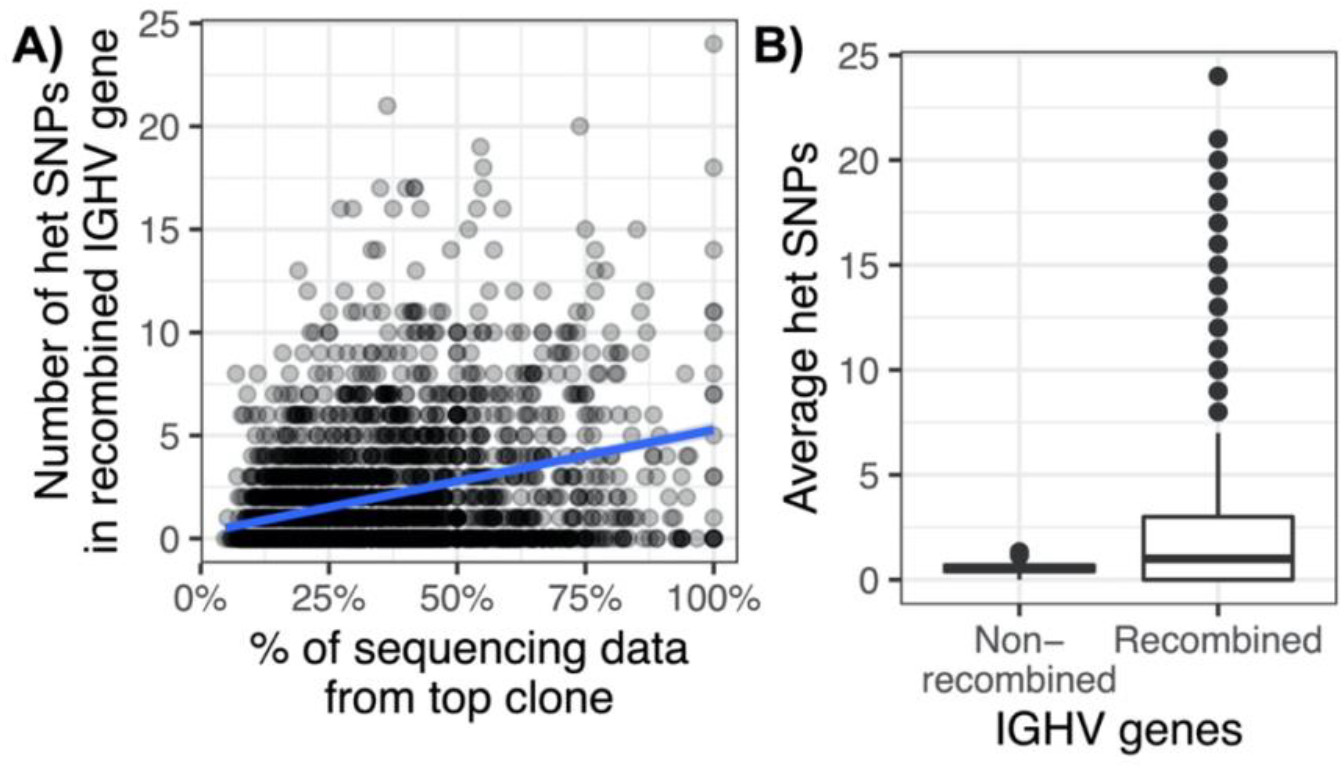
Effect of V(D)J recombination and clonality on heterozygous SNP calling. (A) The number of heterozygous SNP in the V(D)J IGHV selected gene compared to the percentage of sequencing data from dominant clone (B) The average number of heterozygous SNPs between IGHV genes selected for V(D)J recombination and not selected

**Supp Fig 4.**
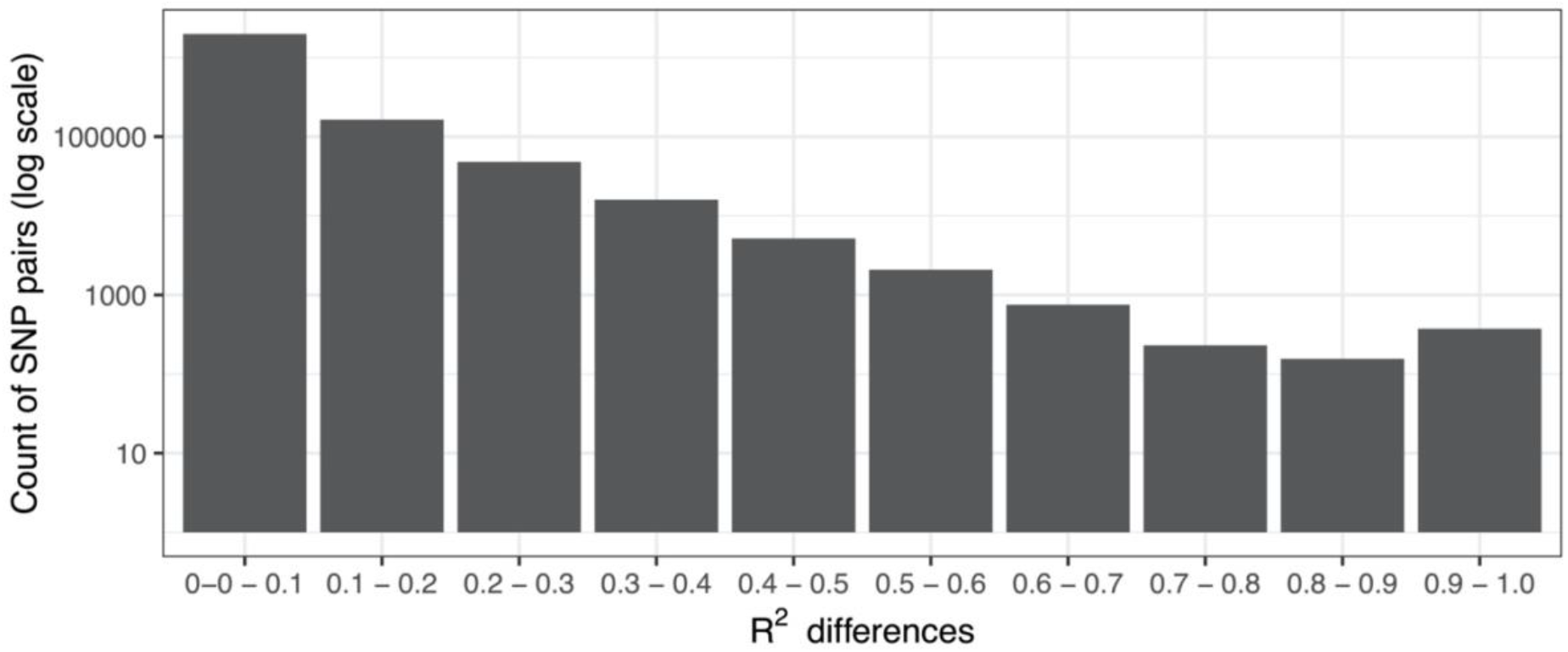
LD difference between African individuals with low clonality (0-25%) and high clonality (75%-100%).

